# Stability of DNA-Methylation Profiles of Biological Aging in Children and Adolescents

**DOI:** 10.1101/2023.10.30.564766

**Authors:** Abby J. deSteiguer, Laurel Raffington, Aditi Sabhlok, Peter Tanksley, Elliot M. Tucker-Drob, K. Paige Harden

**Affiliations:** Department of Psychology, University of Texas at Austin, Austin, TX, USA; Max Planck Research Group Biosocial – Biology, Social Disparities, and Development, Max Planck Institute for Human Development, Berlin, Germany; Population Research Center, The University of Texas at Austin, Austin, TX, USA

**Author notes:** **Address correspondence to:** Abby J. deSteiguer, Department of Psychology, University of Texas at Austin, 1 University Station A8000, Austin, TX 78712,. **Role of Funder/Sponsor:** The funders or sponsors did not participate in the work. Funded by the National Institutes of Health (NIH). **Availability of Data and Material:** Because of the high potential for deductive identification in this special population from a geographically circumscribed area, and the sensitive nature of information collected, data from the Texas Twin Project are not shared with individuals outside of the research team. **Code Availability:** Code will be shared by the first author upon request. **Author Contributions:** Abby J. deSteiguer conceptualized and designed the study, analyzed the data, drafted the manuscript, and critically revised the manuscript; Dr Laurel Raffington conceptualized and designed the study and critically revised the manuscript; Dr K. Paige Harden conceptualized and designed the study, collected the data, drafted the manuscript, and critically revised the manuscript; Dr Aditi Sabhlok collected the data and critically revised the manuscript; Dr Peter Tanksley critically revised the manuscript; Dr Elliot M. Tucker-Drob conceptualized and designed the study, collected the data, and critically revised the manuscript; all authors reviewed the manuscript and approved the final manuscript as submitted and agree to be accountable for all aspects of the work.

## Abstract

**Background and Objectives:** Methylation profile scores (MPSs) index biological aging and aging-related disease in adults and are cross-sectionally associated with social determinants of health in childhood. MPSs thus provide an opportunity to trace how aging-related biology responds to environmental changes in early life. Information regarding the stability of MPSs in early life is currently lacking.

**Method:** We use longitudinal data from children and adolescents ages 8-18 (N = 428, M age = 12.15 years) from the Texas Twin Project. Participants contributed two waves of salivary DNA-methylation data (mean lag = 3.94 years), which were used to construct four MPSs reflecting multi-system physiological decline and mortality risk (PhenoAgeAccel and GrimAgeAccel), pace of biological aging (DunedinPACE), and cognitive function (Epigenetic-*g*). Furthermore, we exploit variation among participants in whether they were exposed to the COVID-19 pandemic during the course of study participation, in order to test how a historical period characterized by environmental disruption might affect children’s aging-related MPSs.

**Results:** All MPSs showed moderate longitudinal stability (test-retest *r*s = 0.42, 0.44, 0.46, 0.51 for PhenoAgeAccel, GrimAgeAccel, and Epigenetic-*g*, and DunedinPACE, respectively). No differences in the stability of MPSs were apparent between those whose second assessment took place after the onset of the COVID-19 pandemic vs. those for whom both assessments took place prior to the pandemic.

**Conclusions:** Aging-related DNA-methylation patterns are less stable in childhood than has been previously observed in adulthood. Further developmental research on the methylome is necessary to understand which environmental perturbations in childhood impact trajectories of biological aging and when children are most sensitive to those impacts.

**Article Summary:** Methylation profiles of biological aging are less stable in childhood than has been previously observed in adulthood.

**What’s Known on This Subject:** Methylation profile scores (MPSs) index biological aging in adults and are cross-sectionally associated with social determinants of health in childhood. Aging-related MPSs in adulthood show very high test-retest stability but data on longitudinal stability of MPSs in childhood is sparse.

**What This Study Adds:** Children’s methylation profiles of biological aging are moderately stable across an approximately four-year period. Methylation profiles are less stable in childhood than in adulthood, suggesting that aging-related biology in childhood might be more responsive to environmental changes than in adulthood.

## Introduction

Childhood social adversity is linked to lifespan risk for aging-related diseases like cardiovascular disease, diabetes, cancer, dementia, and early mortality.^1–5^ Recent advances in - omics technologies have led to the identification of DNA-methylation (DNAm) profiles that index processes of biological aging and aging-related diseases. These DNAm profile scores (MPSs) provide an opportunity to trace how aging-related biology responds to environmental changes in early life, years or even decades before the negative health sequelae of childhood experiences are observable.^6,7^ However, basic information regarding the stability and plasticity of MPSs in early life is currently lacking. Here, we use longitudinal data from a pediatric sample to investigate the early stability of trajectories of aging-related biology as measured by MPSs.

Previous studies in adults have identified systematic patterns of DNAm differences that track a variety of aging-related phenotypes, including (1) biological age as measured by multi-system physiological decline and mortality risk (PhenoAge^8^ and GrimAge^9^), (2) the pace of biological aging across physiological systems assessed longitudinally across midlife (DunedinPACE^10^), and (3) cognitive functions accounting for chronological age (Epigenetic-g^11^).These MPSs are now amongst the strongest predictors of all-cause morbidity, mortality, and physiological decline in adults,^12,13^ offering researchers the opportunity to measure progressive system decline occurring in cells, tissues, and organs. Moreover, MPSs of biological aging, when measured in children, adolescents and young adults, are associated with known social determinants of life course health. Specifically, child victimization, family and neighborhood-level socioeconomic disadvantage in childhood, and belonging to a marginalized racial or ethnic group are associated cross-sectionally with MPSs that index more advanced biological age, a faster pace of aging, and lower cognitive performance in childhood.^14–17^ Together, this research suggests that MPSs might be useful for quantifying, in real time, how early life experiences affect aging-related biology. However, there is currently very little information regarding when aging-related DNAm patterns are most sensitive to environmental change, versus when they are canalized into relatively stable trajectories.

Adults’ aging-related biology shows high *homeorhesis,*^18^ a term that comes from the Greek for “similar flow”: Adults show highly consistent trajectories of DNAm change despite perturbations to their environments. Specifically, aging-related MPSs in middle-adulthood calculated from whole blood are very stable over time, with 16-year test-retest correlations of 0.73, 0.85, and 0.93 for DunedinPACE, PhenoAge, and GrimAge, respectively,^19^ Aging-related biology is often hypothesized to be more plastic and more responsive to environmental perturbations earlier in development,^20–22^ but longitudinal investigations of DNAm in pediatric samples remain rare. Previous work on telomere length has found evidence for high rank-order stability already in childhood (measured at age 11 and 14-year follow-up).^23^ Similarly, some saliva-based MPSs have been found to be fairly stable in adolescence: a salivary MPS of BMI showed a test-retest correlation of 0.63 from age 9 to 15,^24^ while Horvath epigenetic age measured in saliva showed a 2-year test-retest stability of *r* = 0.73 in early adolescence (mean age at baseline = 12.5 years).^25^ Whether other salivary MPSs show more longitudinal instability in childhood and adolescence is unknown, and this information can provide valuable context for understanding when aging-related biology might be most sensitive to environmental intervention.

Stability may be attenuated when there are major changes in the environment with heterogeneous impacts on individuals. In the current study, we examine the longitudinal stability of aging-related MPSs in a sample spanning the ages of 8 to 18. Furthermore, we exploit an exogeneous environmental shock – the onset of the COVID-19 pandemic, which not only exposed millions of children to a novel virus, but also profoundly disrupted their school, family, and peer environments.^26–29^ While pandemic-mitigation policies resulted in a historic decrease in child poverty levels in the United States,^30^ the pandemic may have widened already extant disparities related to other psychosocial factors including isolation, lack of socialization, mental health concerns, disruptions in formal schooling, and material deprivation.^28,29,31–33^ The heterogeneous impact of the pandemic on individuals’ lives may have introduced heterogeneity in experiences relevant for DNAm. A secondary aim of the study, therefore, was to test whether the developmental stability of aging-related MPSs in childhood was affected by exposure to pandemic-related environmental disruptions.

## Methods

### Sample

The Texas Twin Project (TTP) is a population-based study of school-aged twins in Austin and central Texas.^34^ Ethical approval is granted by the University of Texas at Austin Institutional Review Board. Data were collected between 2014 and 2022, with a pause in collection from March 2020 until June 2021. Of N = 428 participants, N = 245 contributed two DNAm samples prior to the onset of the COVID-19 pandemic (Pre-COVID), and N = 183 contributed one sample prior to and one sample after the onset of the pandemic (Peri-COVID). Sample demographics are in **Table 1**, and overlap between wave, batch, and COVID-19 group is shown in **Table S1.** Data acquisition by date of assessment and by chronological age is illustrated in **Figure 1**.

**Figure 1.**
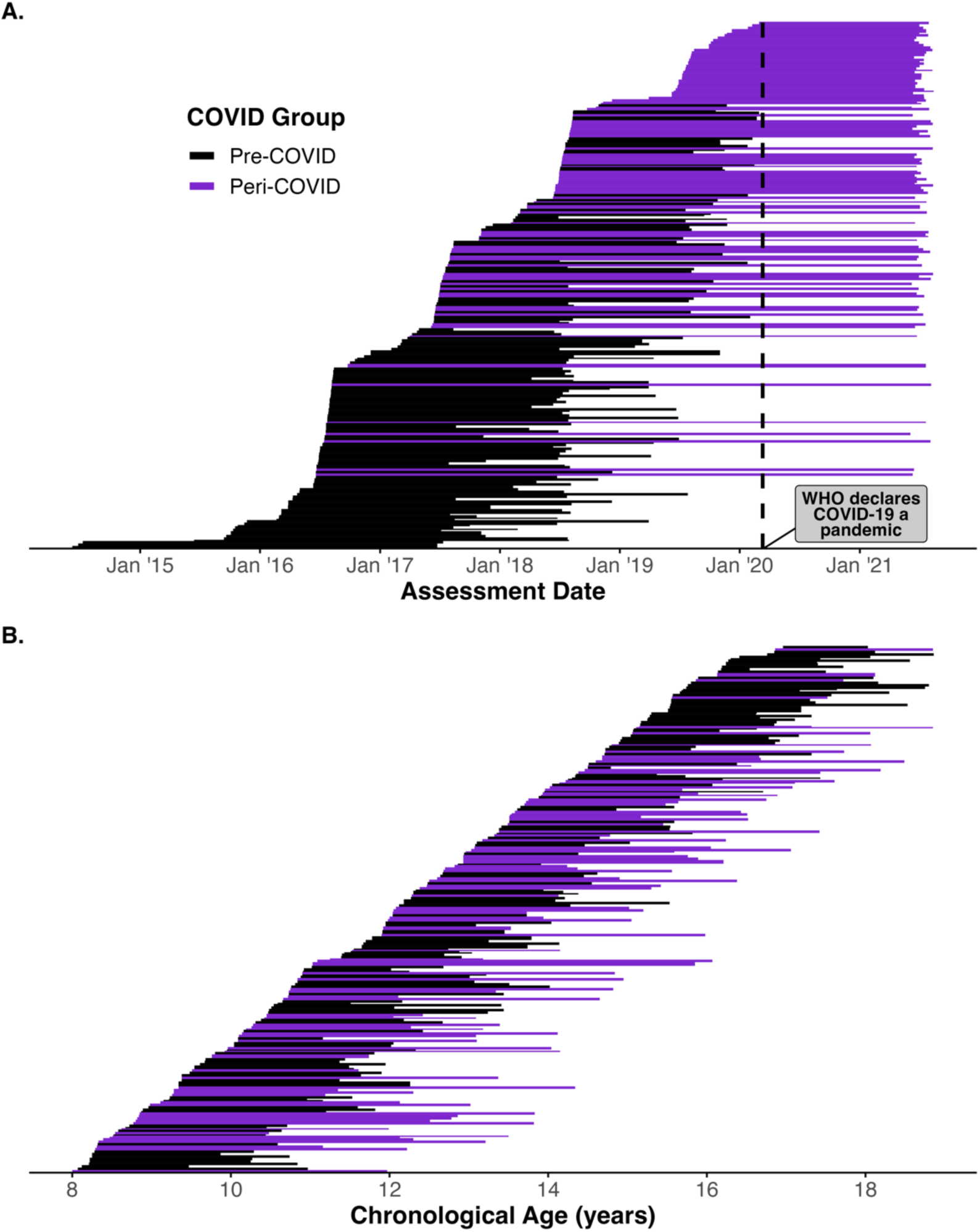
Acquisition of Longitudinal DNAm Data by (A) Date of Assessment and by (B) Participant Age. Pre-COVID group (Black) includes participants with two waves of DNAm data prior to the onset of the COVID-19 pandemic. Peri-COVID group (Purple) includes participants with one wave of data prior and one wave after the onset of the COVID-19 pandemic. The large increases in sample size over short periods of time are concentrated in summer months when participant sessions took place five to seven days per week, in comparison to during the school year when sessions primarily took place on weekends. No sessions were conducted during the first year of the COVID-19 pandemic, creating a longer average lag in assessment for those in the Peri-COVID group.

**Table 1.**
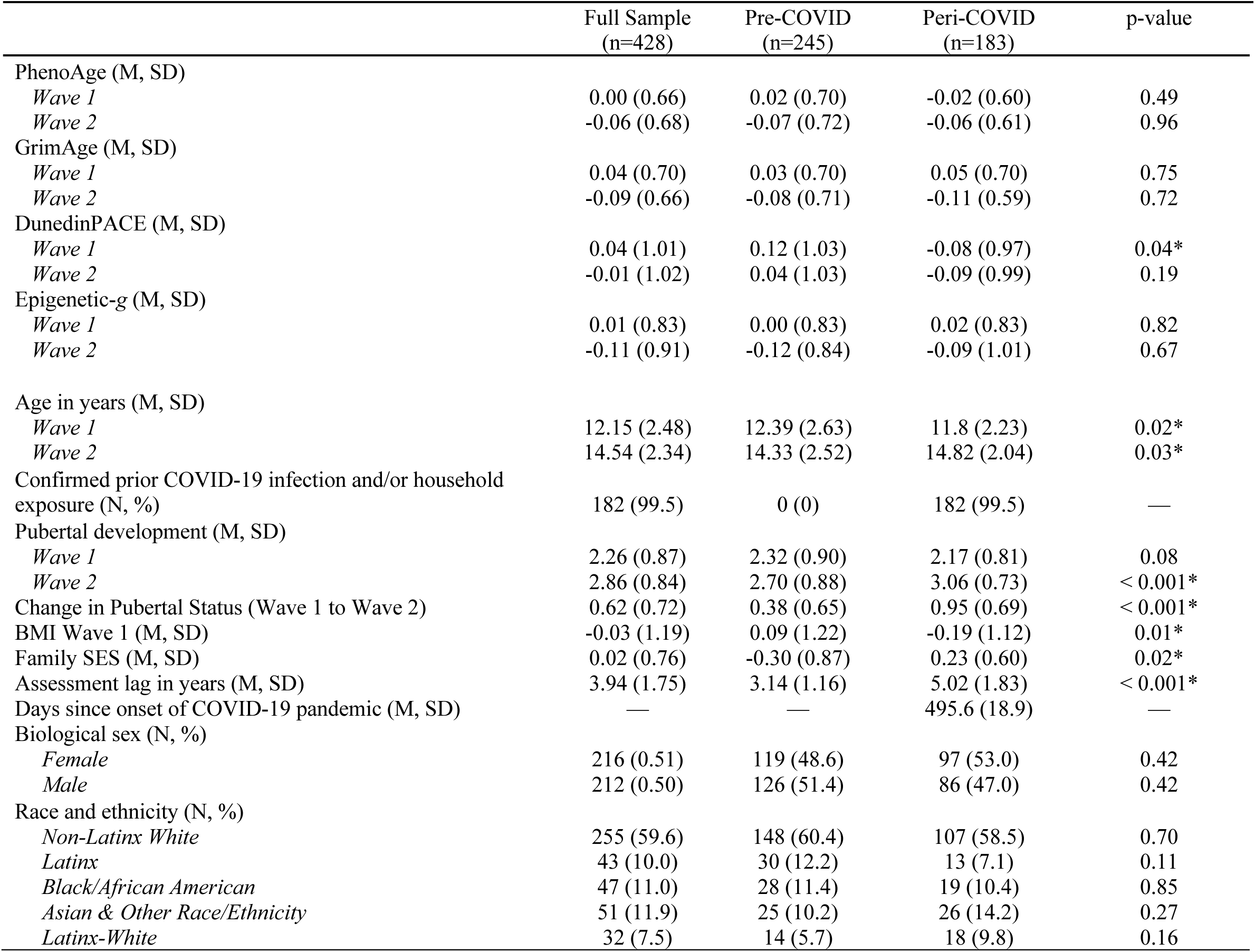
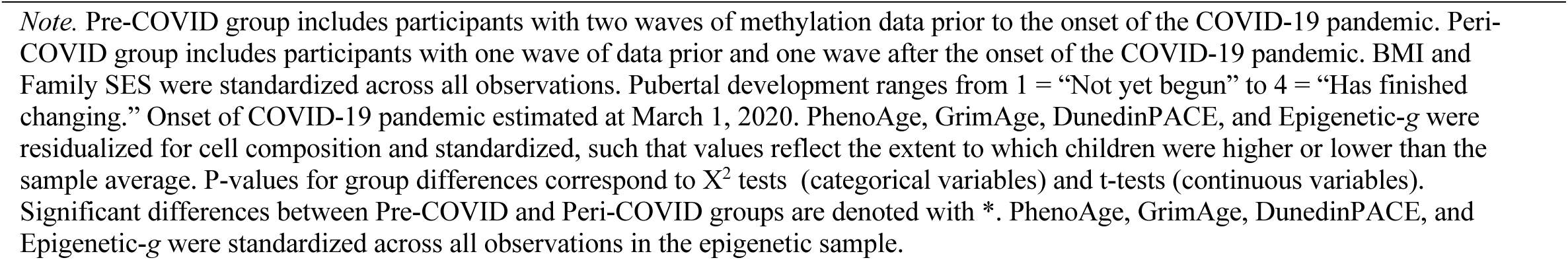
Descriptive Statistics of Study Measures.

### Measures

#### DNAm preprocessing

Saliva samples (2 mL) were collected using Oragene kits (DNA Genotek, Ottawa, ON, Canada). DNA extraction and methylation profiling was conducted by Edinburgh Clinical Research Facility (UK). The Infinium MethylationEPIC BeadChip kit (Illumina, Inc., San Diego, CA) was used to assess methylation levels at 850,000 methylation sites. DNAm preprocessing was primarily conducted with the ‘minfi’ package in R.^35^ Within-array normalization was performed to address array background correction, red/green dye bias, and probe type I/II correction.^36^ Noob preprocessing as implemented by minfi’s “preprocessNoob”^37^ is a background correction and dye-bias equalization method that has similar within-array normalization effects on the data as probe type correction methods such as BMIQ.^38^

CpG probes with detection p > 0.01 and fewer than 3 beads in more than 1% of the samples and probes in cross-reactive regions were excluded.^39^ Samples were excluded if (1) they showed low intensity probes as indicated by the log of average methylation <9 and their detection p is > 0.01 in >10% of their probes, (2) their self-reported and methylation-estimated sex mismatched, and (3) their self-reported and DNA-estimated sex mismatched. Cell composition of immune and epithelial cell types (i.e., CD4+ T-cell, natural killer cells, neutrophilseosinophils, B cells, monocytes, CD8+ T-cell, and granulocytes) was estimated using a child saliva reference panel implemented in the R package “BeadSorted.Saliva.EPIC” within “ewastools”.^40^ Surrogate variable analysis was used to correct methylation values for batch effects using the “combat” function in the SVA package,^41^ which is important for longitudinal analyses. Longitudinal wave was not completely confounded with each batch and there were 20 technical repeats to assess reliability.

#### MPSs

MPSs for PhenoAge, GrimAge, DunedinPACE and Epigenetic-*g* were computed using algorithms from previous epigenome-wide association studies (EWAS) and elastic net analyses (see **Table 2**; DunedinPACE and epigenetic-*g* were preregistered outcomes. PhenoAge and GrimAge were added subsequently due to their relevance for health and mortality and their strong associations with age). All MPSs were residualized for array, slide, batch, cell composition and then standardized in the pooled sample (Mean = 0, SD = 1) to ease interpretation. Analysis of 20 technical replicates in our sample suggests that measurement reliability of the MPSs analyzed here (residualized for technical artifacts and cell composition) is good (intraclass correlation coefficients for PhenoAge = 0.79, GrimAge = 0.85, DunedinPACE = 0.78, epigenetic-*g =* 0.73). See **Table 2** for a description of covariate measures.

**Table 2.** Description of Measures.

| Measure | Description |
| --- | --- |
| PhenoAge<sup>8</sup> | PhenoAge was calculated using the algorithm available at https://github.com/MorganLevineLab/PC-Clocks. PhenoAge is an age estimate, with greater scores indicating greater biological age. The PhenoAge algorithm was originally trained on nine clinical biomarkers (e.g., albumin, creatinine, C-reactive protein, and white blood cell count) and chronological age in adults ages 21 to 100. |
| GrimAge<sup>9</sup> | GrimAge was calculated using the algorithm available at https://github.com/MorganLevineLab/PC-Clocks. GrimAge is an age estimate, with greater scores indicating greater biological age. GrimAge was trained on DNAm-based smoking pack-years, age, sex, and seven DNAm-based surrogate markers of plasma proteins. |
| DunedinPACE pace of aging<sup>10</sup> | DunedinPACE was calculated as a composite phenotype calculated using a published algorithm available at https://github.com/danbelsky/DunedinPACE. Scores indicate years of physiological decline experience per calendar year. The expected value of DunedinPACE in midlife adults is 1. Values >1 indicate accelerated aging, while values <1 indicate slowed aging. DunedinPACE measures the rate at which aging occurs and was derived from analysis of longitudinal changes at 19 biomarkers (e.g., body mass index, glycated hemoglobin, total cholesterol, cardiorespiratory fitness, and tooth decay) measured at 26, 32, 38, and 45 years of age in the Dunedin Study. |
| Epigenetic-g<sup>11</sup> | Epigenetic-g was calculated using the algorithm available at https://gitlab.com/danielmccartney/ewas_of_cognitive_funct. Prior to computation, methylation values were scaled within each CpG site (mean = 0, SD = 1). Higher scores represent greater cognitive function. A higher score indicates a profile that more closely resembles the profile of adults from the Generation Scotland Study who performed higher on tests of general function, including logical memory, digit symbol test score, verbal fluency, and vocabulary. Epigenetic-g was computed using a blood-based algorithm from an epigenome-wide association study of general cognitive functions in adults ages 18 to 93. |
| Assessment lag | Lag between first and second assessments was calculated in years and centered at the pooled sample mean lag (3.94 years); see **Figure 1** for visualization of assessment lag. |
| DNAm-Smoke | A poly-DNA-methylation measure of tobacco exposure was derived by summing the product of the weight and the individual beta estimate for each individual at each CpG site significantly associated with cigarette smoking in a discovery EWAS<sup>62</sup>. |
| Family-level Socioeconomic Status (SES) | A family-level SES index score was computed as a composite of standardized, log transformed parent-reported household income and average, standardized parental education level. |
| COVID-19-related Socioeconomic Stress | Parent-reported socioeconomic stressors unique to the pandemic were used to compute a dummy-coded variable based on loss of employment income and/or received or applied for unemployment insurance, with a score of 1 indicating experiencing one or both of these stressors. |
| Body Mass Index (BMI) | BMI at Wave 1 was determined from measurements of height and weight and was transformed to gender and age-normed z-scores according to the US Centers for Disease Control and Prevention method (https://www.cdc.gov/growthcharts/percentile_data_files.htm). Height and weight were measured in-laboratory. |
| Pubertal Development | Self-reported pubertal development was measured with the Pubertal Development Scale<sup>63</sup>, which shows good reliability and validity, to assess pubertal development across five sex-specific domains. Pubertal status score was measured computing the average response (1 = “Not yet begun” to 4 = “Has finished changing”) across items, and was residualized for age, gender, and age by gender. |

### Statistical Analyses

Our analysis plan was pre-registered in Open Science Framework (OSF; https://osf.io/mfybr/). We first assessed how each MPS was associated with chronological age. In all subsequent analyses, MPSs were residualized for chronological age, thus MPS values reflect the extent to which children were higher or lower than other children their age. For each age-residualized MPS, we conducted a series of longitudinal models regressing Wave 2 MPS on Wave MPS and on other predictors and covariates of interest. We fit regressions using linear mixed models implemented with the *lme4* R package.^42^ All models included a random intercept, representing the family-level intercept of the dependent variable, to correct for non-independence of twins within families. Standardized betas and 95% confidence intervals are reported for each model. A *p*-value of less than 0.05 indicated a statistically significant effect of the predictor variable(s) on the dependent variables. Within each set of models, we controlled for multiple testing using the Benjamini–Hochberg false discovery rate (FDR) method (using a family of four tests, representing each of the four MPSs under study),^43^ Nominal and FDR-adjusted *p*-values are denoted in each table.

## Results

### Methylation Profile Scores Show Variable Patterns of Age-Related Mean Change

All MPSs were significantly correlated with age (PhenoAge: *r* = 0.69, 95% CI [0.65, 0.72]; GrimAge: *r* = 0.68, 95% CI [0.65, 0.72]; DunedinPACE: *r* = 0.08, 95% CI [0.02, 0.15]; Epigenetic-*g*: *r* = 0.35, 95% CI [0.29, 0.41]. That is, all MPSs were higher in older than in younger children, with the strongest age differences evident for MPSs that included age as a training variable (PhenoAge, GrimAge), rather than trained on phenotypic differences between age-matched participants (DunedinPACE, Epigenetic-*g*). To formally test within-person age-related change in MPSs, we regressed each MPS on both between-person age *differences* (an individual’s mean age across two waves) and within-person age *changes* (the deviation between an individual’s age at each wave from their mean age across waves). Results show that there was significant within-person age-related change in GrimAge, which increased by 0.19 per year (b = 0.19, 95% CI [0.17, 0.22], t(850) = 14.30, p < .001) and in PhenoAge (b = 0.22, 95% CI [0.19, 0.25], t(850) = 16.31, p < .001). There was also within-person age-related change in Epigenetic-g (b = 0.10, 95% CI [0.07, 0.14], t(850) = 6.00, p < .001), but not in DunedinPACE (beta = 0.02, 95% CI [-0.02, 0.05], t(850) = 0.88, p = 0.381). Plotting the individual age-related trajectories from Wave 1 to Wave 2 (**Figure 2**) suggests that there was considerable variability around the mean age-related trends.

**Figure 2.**
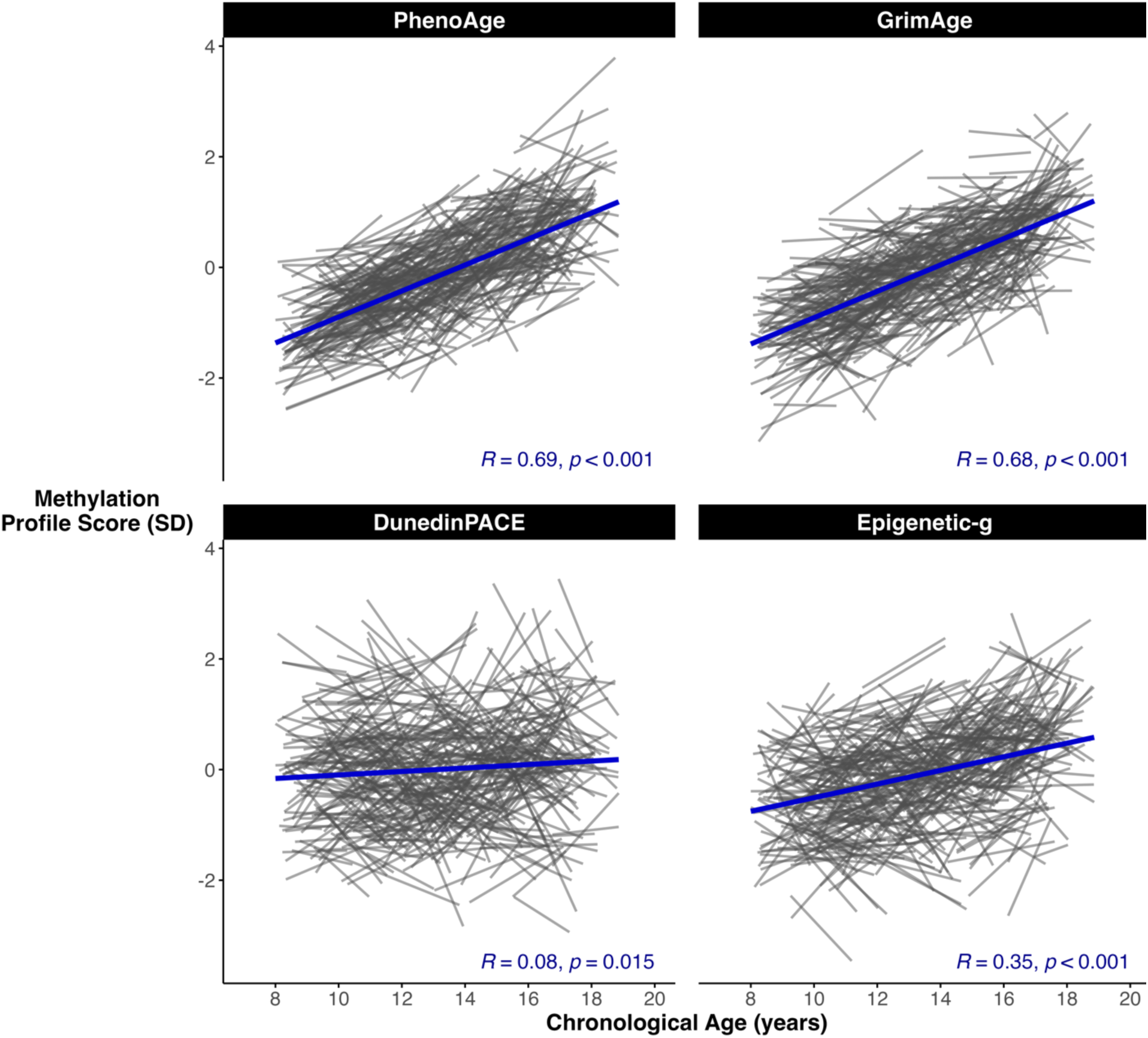
Age-Related Changes in Methylation Profiles Scores (MPSs) in Childhood. Each line represents one individual’s MPS trajectory from Wave 1 to Wave 2. PhenoAge and GrimAge are measures of biological age and mortality risk, with greater scores indicating greater biological age. DunedinPACE is a measure of pace of biological aging, with scores indicating years of physiological decline per calendar year. Epigenetic-g is a measure of cognitive function in adults accounting for chronological age, with higher scores reflective of higher general cognitive function.

### MPSs Exhibit Moderate Rank-Order Stability in Children and Adolescents

All subsequent analyses residualized each MPS for the effects of chronological age, so that estimates of rank-order stability are not biased by age-related differences within our age-heterogeneous sample. The resulting “acceleration” variables represent how a child’s MPS compared to other children the same age. For each MPS, the two waves of data were moderately correlated (PhenoAgeAccel: Pearson’s *r* = 0.42, 95% CI 0.34 to 0.49; GrimAgeAccel: Pearson’s *r* = 0.44, 95% CI 0.36 to 0.51; DunedinPACE: Pearson’s *r* = 0.51, 95% CI 0.43 to 0.57; Epigenetic-*g*: Pearson’s *r* = 0.46, 95% CI 0.38 to 0.53). These results suggest that child MPSs show considerably less stable trajectories of aging-related DNAm than do adult MPSs.^19^ **Figure 3** compares these test-retest correlations to the developmental stability of other aspects of health and development. Relative to other phenotypes in childhood,^52–58^ MPSs show about as much longitudinal (in)stability as child personality, which is considerably less stable than anthropometric or cognitive measurements.

**Figure 3.**
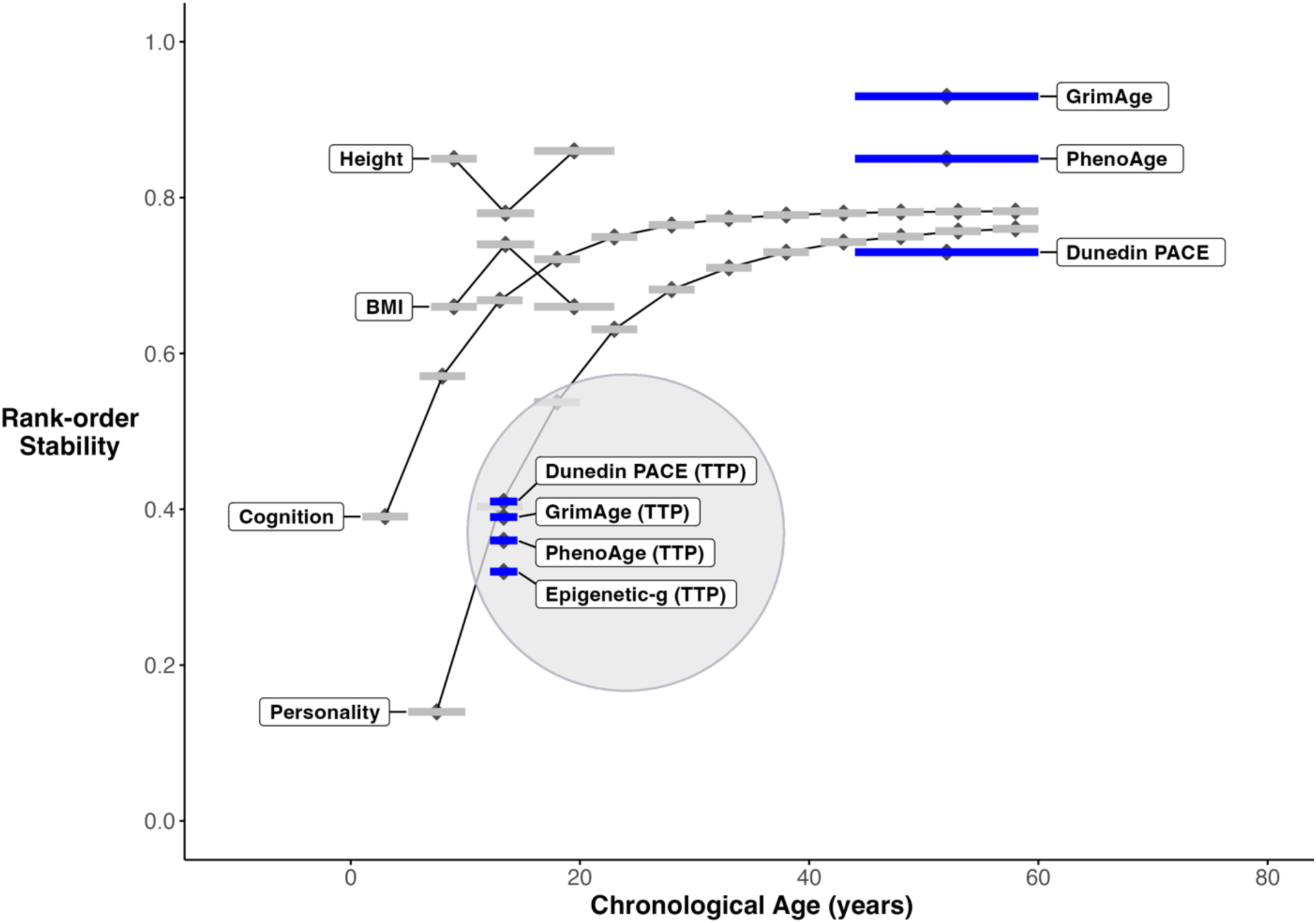
Rank-order stability (test-retest correlations) for child/adolescent methylation profile scores of accelerated biological age, pace of aging, and cognitive performance (∼ 4-year lag) in comparison to other aspects of child health and development. Bayer et al. 2011^44^; Height (4- to 7-year lag; estimates averaged for males and females): Power et al., 1997, BMI (4- to 7-year lag; estimates averaged for males and females): Power et al., 1997^45^; Personality (5-year lag): Bleidorn et al., 2022^46^; Cognition (6-year lag): Tucker-Drob & Briley^47^, 2014; Adult MPSs (16-year lag):Reed et al., 2022.^19^

To formally test the rank-order stability of MPSs, we first regressed Wave 2 MPS on Wave 1 MPS (**Table 3, Model 1**), and then statistically controlled for demographic covariates. Results were generally unchanged upon addition of covariates, with βs ranging from 0.37 to 0.42 (**Table 3, Model 2**). The previous analyses assess stability at the mean lag in assessment of 3.94 years (by including a term of lag centered at the mean lag). Upon inclusion of a Lag*Wave 1 interaction term, which allows for an exploration of the effect of stability at different lags in assessment, main effect estimates of stability are nearly identical to the model without an interaction term (PhenoAgeAccel: β = 0.37, 95% CI 0.28, 0.46; GrimAgeAccel: β = 0.39, 95% CI 0.29, 0.49; DunedinPACE: β = 0.41, 95% CI 0.33, 0.50; Epigentic-*g:* β = 0.42, 95% CI 0.33, 0.52). The interaction terms were non-significant (PhenoAgeAccel: β = 0.02, 95% CI -0.06, 0.11; GrimAgeAccel: β = 0.02, 95% CI -0.07, 0.10; DunedinPACE: β = -0.05, 95% CI -0.13, 0.04; Epigentic-*g:* β = 0.04, 95% CI -0.06, 0.13), suggesting that stability did not increase or decrease over different lags. Additionally, stability did not vary over different ages at first assessment (**Supplemental Table 2**). Means and variances of MPSs at Wave 1 and Wave 2 are reported in **Table 1**.

**Table 3.** Regression of MPSs at Wave 2 on Wave 1.

|  | PhenoAgeAccel<br>(Wave 2) |  | GrimAgeAccel (Wave 2) |  | Dunedin PACE (Wave 2) |  | Epigenetic-g (Wave 2) |  |
| --- | --- | --- | --- | --- | --- | --- | --- | --- |
| *Model 1 (n=428)* | *std. Beta* | *95% CI* | *std. Beta* | *95% CI* | *std. Beta* | *95% CI* | *std. Beta* | *95% CI* |
| MPS Wave 1 | **0.38\*** | 0.29, 0.47 | **0.41\*** | 0.32, 0.50 | **0.45\*** | 0.37, 0.53 | **0.42\*** | 0.33, 0.51 |
| *Model 2 (n=428)* | *std. Beta* | *95% CI* | *std. Beta* | *95% CI* | *std. Beta* | *95% CI* | *std. Beta* | *95% CI* |
| MPS Wave 1 | **0.37\*** | 0.28, 0.46 | **0.39\*** | 0.29, 0.49 | **0.41\*** | 0.33, 0.50 | **0.42\*** | 0.33, 0.51 |
| Group (Peri-COVID) | -0.07 | -0.31, 0.16 | -0.19 | -0.42, 0.04 | -0.06 | -0.27, 0.16 | -0.06 | -0.29, 0.18 |
| Assessment Lag | 0.08 | -0.04, 0.20 | **0.14** | 0.02, 0.25 | -0.01 | -0.12, 0.09 | 0.09 | -0.03, 0.20 |
| Sex (Male) | **-0.42\*** | -0.60, -0.25 | 0.09 | -0.10, 0.28 | **-0.32\*** | -0.48, -0.16 | -0.01 | -0.19, 0.17 |
| Latinx | 0.02 | -0.08, 0.12 | 0.07 | -0.03, 0.16 | **0.14\*** | 0.05, 0.24 | -0.02 | -0.12, 0.08 |
| BlackPlus | **0.16\*** | 0.06, 0.26 | **0.12\*** | 0.02, 0.22 | **0.11\*** | 0.02, 0.21 | -0.06 | -0.16, 0.04 |
| AsianPlusOther | 0.02 | -0.08, 0.12 | -0.01 | -0.11, 0.09 | 0.04 | -0.05, 0.13 | -0.03 | -0.13, 0.06 |
| LatinxWhite | 0.01 | -0.09, 0.12 | 0.02 | -0.07, 0.12 | **0.09** | 0.00, 0.18 | 0.01 | -0.09, 0.11 |
| DNAm-Smoke | -0.06 | -0.15, 0.03 | 0.02 | -0.08, 0.12 | **0.10** | 0.02, 0.19 | -0.09 | -0.18, 0.00 |
| *Model 3 (N=251)* | *std. Beta* | *95% CI* | *std. Beta* | *95% CI* | *std. Beta* | *95% CI* | *std. Beta* | *95% CI* |
| MPS Wave 1 | **0.36\*** | 0.23, 0.48 | **0.39\*** | 0.25, 0.52 | **0.44\*** | 0.32, 0.55 | **0.43\*** | 0.31, 0.54 |
| Group (Peri-COVID) | -0.00 | -0.34, 0.33 | -0.14 | -0.45, 0.18 | -0.14 | -0.45, 0.17 | -0.19 | -0.51, 0.13 |
| Assessment Lag | 0.11 | -0.05, 0.27 | 0.14 | -0.02, 0.29 | 0.07 | -0.08, 0.21 | 0.09 | -0.06, 0.25 |
| Sex (Male) | **-0.38\*** | -0.62, -0.13 | 0.20 | -0.06, 0.47 | **-0.43\*** | -0.65, -0.20 | -0.06 | -0.30, 0.18 |
| Latinx | 0.00 | -0.14, 0.14 | -0.05 | -0.18, 0.09 | 0.12 | -0.01, 0.25 | -0.02 | -0.16, 0.11 |
| BlackPlus | 0.10 | -0.03, 0.24 | 0.07 | -0.06, 0.20 | 0.12 | -0.01, 0.25 | -0.11 | -0.25, 0.02 |
| AsianPlusOther | 0.02 | -0.12, 0.15 | -0.01 | -0.13, 0.12 | 0.02 | -0.11, 0.14 | -0.05 | -0.18, 0.08 |
| LatinxWhite | -0.04 | -0.17, 0.10 | -0.04 | -0.16, 0.09 | 0.09 | -0.03, 0.21 | -0.02 | -0.15, 0.10 |
| DNAm-Smoke | 0.01 | -0.12, 0.14 | 0.07 | -0.06, 0.20 | 0.08 | -0.03, 0.20 | -0.01 | -0.13, 0.11 |
| Family SES (Wave 2) | 0.02 | -0.13, 0.17 | -0.09 | -0.23, 0.05 | -0.06 | -0.20, 0.08 | 0.09 | -0.05, 0.23 |
| BMI (Wave 1) | 0.05 | -0.08, 0.17 | 0.01 | -0.11, 0.13 | -0.00 | -0.12, 0.11 | -0.02 | -0.14, 0.09 |
| PDS (Wave 1) | 0.02 | -0.14, 0.19 | 0.02 | -0.14, 0.18 | 0.01 | -0.14, 0.16 | 0.05 | -0.11, 0.21 |
| ΔPDS | -0.04 | -0.20, 0.13 | -0.06 | -0.22, 0.10 | 0.12 | -0.04, 0.27 | 0.01 | -0.15, 0.17 |

Note. MPS = DNA-methylation profile score. CI = Confidence Interval. Nominally significant parameters at $p < 0.05$ are in bold; parameters significant after FDR correction are denoted with \*. All MPSs were age-residualized.

There were statistically significant sex differences in Wave 2 MPSs. Males exhibited lower PhenoAgeAccel (β = -0.42, 95% CI -0.60 to -0.25 and age-residualized DunedinPACE (β = -0.32, 95% CI -0.48 to -0.16); **Table 3, Model 2**). There were also significant race/ethnic differences in PhenoAgeAccel and age-residualized DunedinPACE. Participants identifying as Latinx exhibited faster pace of aging as indicated by age-residualized DunedinPACE compared to those identifying as White (Latinx: β = 0.14, 95% CI 0.05 to 0.24). Participants who identified as Black exhibited a faster pace of aging and more advanced biological age as indicated by age-residualized DunedinPACE (β = 0.11, 95% CI 0.02 to 0.21), PhenoAgeAccel (β = 0.16, 95% CI 0.06 to 0.26), and GrimAgeAccel (β = 0.12, 95% CI 0.02 to 0.22; **Table 3, Model 2)** compared to those identifying as White. That these race/ethnic differences were apparent for Wave 2 MPSs even after controlling for Wave 1 MPSs suggests that racial/ethnic disparities are widening over development.

We then controlled for developmental covariates including BMI, pubertal development, and change in pubertal development (pre-registered). We additionally adjusted for family SES due to the significant difference in this variable between groups (the Peri-COVID group had a higher overall SES, consistent with the hypothesis that the effects of socioeconomic hardship on study participation were heightened during the pandemic). Results were unchanged (**Table 3, Model 3).**

To assess whether rank-order stability is socially stratified, we regressed Wave 2 MPS on Wave 1 MPS, SES covariates (Family SES and COVID-19 related socioeconomic stress), and the interactions between Wave 1 DNAm and each SES covariate. There were no statistically significant main effects or interaction effects for any MPS (**Table 5, Models 1, 2 & 3**).

### Rank-Order Stabilities of MPSs Do Not Differ as a Function of COVID-19

To assess differences in rank-order stability between the Pre-COVID and Peri-COVID groups (illustrated in **Figure 4**), we regressed Wave 2 MPS on Wave 1 MPS, COVID-19 group, and on their interaction (**Table 4**). Neither the main effect of COVID-19 group nor its interaction effect with Wave 1 MPS were significantly different from zero for any MPS, indicating that there were no significant differences by COVID-19 group in mean-level change (represented by the main effect of COVID-19 group) or rank-order stability (represented by the interaction effect of COVID-group and Wave 1 DNAm). This is illustrated in **Figure 4** by the similarity of slopes across groups (representing rank-order stability) and the similar position of regression line midpoints relative to the y-axis across groups (representing mean-level change). Results were unchanged upon the addition of demographic and developmental covariates (**Table 4, Models 2 & 3**).

**Figure 4.**
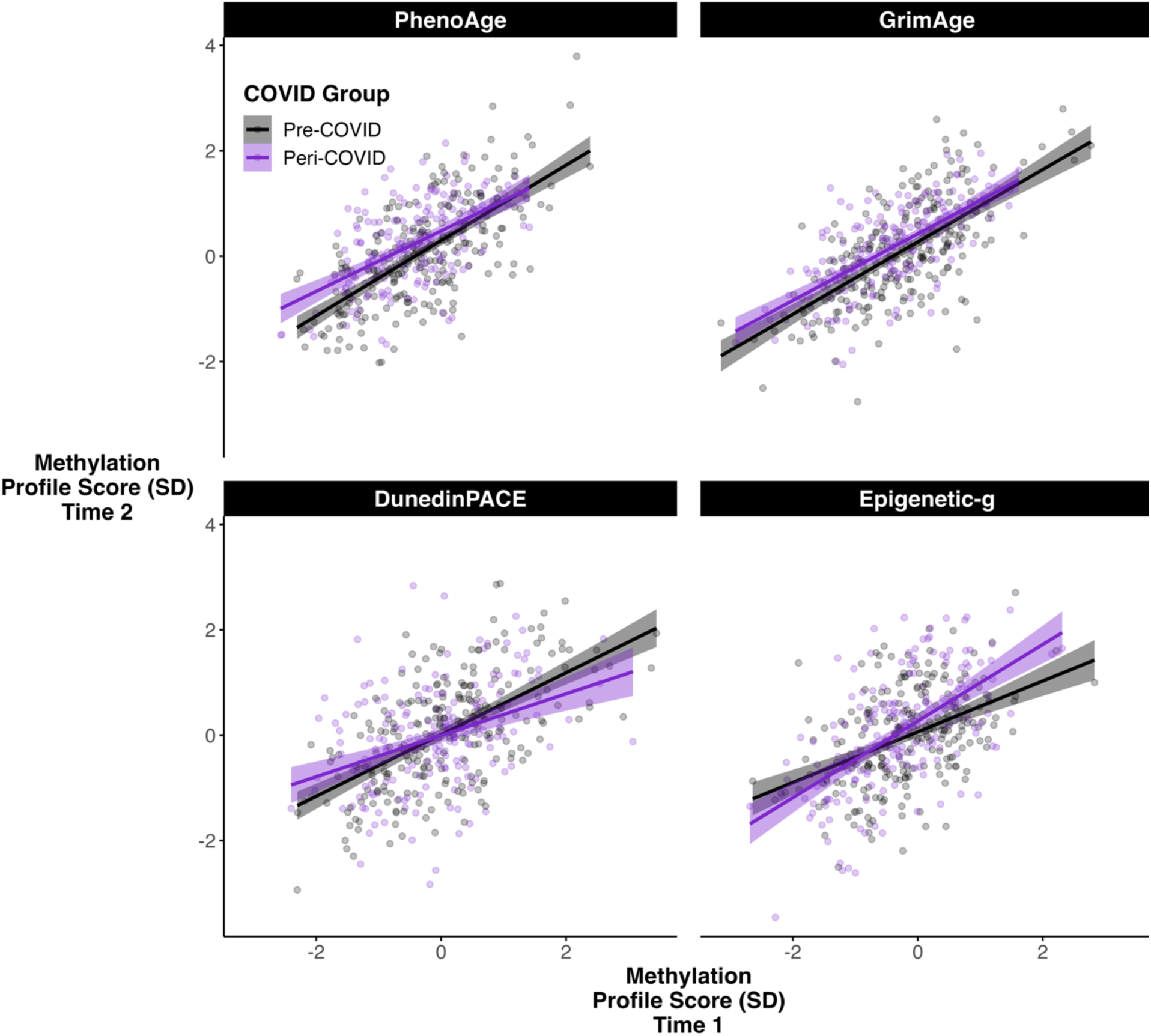
Rank-Order Stability of Age-Residualized Methylation Profiles Scores (MPSs). PhenoAge and GrimAge Acceleration are measures of more advanced biological age, with greater scores indicating older biological age compared to same-aged peers. DunedinPACE is a measure of pace of biological aging, with scores indicating a faster pace of physiological decline per calendar year. Epigenetic-g is a measure of cognitive function, with higher scores reflective of higher general cognitive function. All MPSs were residualized for cell composition and standardized, such that values reflect the extent to which children were higher or lower than other children their age.

**Table 4.** Regression of Wave 2 MPSs on the Interaction between Wave 1 MPSs and COVID-19 Group.

|  | PhenoAgeAccel<br>(Wave 2) |  | GrimAgeAccel (Wave 2) |  | Dunedin PACE (Wave 2) |  | Epigenetic-g (Wave 2) |  |
| --- | --- | --- | --- | --- | --- | --- | --- | --- |
| *Model 1 (n=428)* | *std. Beta* | *95% CI* | *std. Beta* | *95% CI* | *std. Beta* | *95% CI* | *std. Beta* | *95% CI* |
| MPS Wave 1 | **0.38\*** | 0.26, 0.49 | **0.37\*** | 0.25, 0.48 | **0.51\*** | 0.40, 0.62 | **0.36\*** | 0.25, 0.48 |
| Group (Peri-COVID) | 0.02 | -0.18, 0.22 | -0.06 | -0.25, 0.14 | -0.04 | -0.23, 0.14 | 0.03 | -0.16, 0.22 |
| MPS Wave 1 * Group<br>(Peri-COVID) | 0.01 | -0.17, 0.20 | 0.10 | -0.08, 0.28 | -0.15 | -0.32, 0.02 | 0.14 | -0.04, 0.32 |
| *Model 2 (n=428)* | *std. Beta* | *95% CI* | *std. Beta* | *95% CI* | *std. Beta* | *95% CI* | *std. Beta* | *95% CI* |
| MPS Wave 1 | **0.36\*** | 0.25, 0.47 | **0.36\*** | 0.23, 0.49 | **0.46\*** | 0.35, 0.57 | **0.36\*** | 0.24, 0.48 |
| DNAm Wave 1*Group<br>(Peri-COVID) | 0.02 | -0.16, 0.21 | 0.07 | -0.10, 0.25 | -0.11 | -0.28, 0.05 | 0.15 | -0.03, 0.32 |
| Group (Peri-COVID) | -0.07 | -0.31, 0.16 | -0.19 | -0.42, 0.04 | -0.05 | -0.27, 0.16 | -0.05 | -0.28, 0.18 |
| Assessment Lag | 0.08 | -0.04, 0.20 | **0.14** | 0.02, 0.25 | -0.02 | -0.13, 0.09 | 0.08 | -0.03, 0.20 |
| Sex (Male) | **-0.42\*** | -0.60, -0.25 | 0.08 | -0.11, 0.27 | **-0.31\*** | -0.47, -0.14 | 0.00 | -0.18, 0.17 |
| Latinx | 0.02 | -0.08, 0.12 | 0.07 | -0.03, 0.16 | **0.14\*** | 0.05, 0.23 | -0.03 | -0.12, 0.07 |
| BlackPlus | **0.16\*** | 0.06, 0.26 | **0.12\*** | 0.02, 0.21 | **0.11\*** | 0.02, 0.20 | -0.07 | -0.17, 0.03 |
| AsianPlusOther | 0.02 | -0.08, 0.12 | -0.01 | -0.11, 0.09 | 0.04 | -0.05, 0.13 | -0.03 | -0.13, 0.07 |
| LatinxWhite | 0.01 | -0.09, 0.12 | 0.02 | -0.08, 0.12 | **0.09** | 0.00, 0.18 | 0.02 | -0.08, 0.11 |
| DNAm-Smoke | -0.06 | -0.15, 0.03 | 0.02 | -0.08, 0.12 | **0.10** | 0.02, 0.19 | -0.09 | -0.18, 0.00 |
| *Model 3 (N=251)* | *std. Beta* | *95% CI* | *std. Beta* | *95% CI* | *std. Beta* | *95% CI* | *std. Beta* | *95% CI* |
| MPS Wave 1 | **0.37\*** | 0.17, 0.56 | **0.33\*** | 0.13, 0.52 | **0.61\*** | 0.44, 0.79 | **0.36\*** | 0.18, 0.55 |
| Group (Peri-COVID) | 0.00 | -0.34, 0.34 | -0.14 | -0.45, 0.18 | -0.13 | -0.44, 0.17 | -0.18 | -0.50, 0.14 |
| MPS Wave 1* Group (Peri-COVID) | -0.01 | -0.26, 0.24 | 0.10 | -0.14, 0.34 | **-0.28** | -0.50, -0.06 | 0.10 | -0.13, 0.34 |
| Assessment Lag | 0.11 | -0.06, 0.27 | 0.14 | -0.01, 0.29 | 0.04 | -0.11, 0.19 | 0.09 | -0.07, 0.24 |
| Sex (Male) | **-0.38\*** | -0.63, -0.13 | 0.19 | -0.07, 0.46 | **-0.41\*** | -0.63, -0.18 | -0.06 | -0.30, 0.18 |
| Latinx | 0.00 | -0.14, 0.15 | -0.05 | -0.19, 0.08 | **0.13** | 0.00, 0.26 | -0.03 | -0.16, 0.11 |
| BlackPlus | 0.11 | -0.03, 0.24 | 0.06 | -0.07, 0.19 | 0.12 | -0.01, 0.24 | -0.12 | -0.25, 0.02 |
| AsianPlusOther | 0.02 | -0.12, 0.15 | -0.01 | -0.14, 0.12 | 0.01 | -0.11, 0.13 | -0.05 | -0.18, 0.08 |
| LatinxWhite | -0.04 | -0.17, 0.10 | -0.04 | -0.17, 0.09 | 0.10 | -0.02, 0.22 | -0.02 | -0.15, 0.11 |
| DNAm-Smoke | 0.01 | -0.12, 0.14 | 0.07 | -0.06, 0.21 | 0.07 | -0.04, 0.19 | -0.01 | -0.13, 0.11 |
| Family SES | 0.02 | -0.13, 0.17 | -0.09 | -0.23, 0.05 | -0.04 | -0.17, 0.10 | 0.09 | -0.05, 0.23 |
| BMI (Wave 1) | 0.05 | -0.08, 0.17 | 0.02 | -0.10, 0.14 | -0.02 | -0.14, 0.09 | -0.02 | -0.14, 0.10 |
| PDS (Wave 1) | 0.02 | -0.14, 0.19 | 0.02 | -0.14, 0.18 | 0.03 | -0.12, 0.18 | 0.05 | -0.11, 0.21 |
| ΔPDS | -0.04 | -0.20, 0.13 | -0.06 | -0.22, 0.10 | 0.14 | -0.01, 0.29 | 0.00 | -0.16, 0.16 |

Note. MPS = DNA-methylation profile score. CI = Confidence Interval. Nominally significant parameters at $p < 0.05$ are in bold; parameters significant after FDR correction are denoted with \*. All MPSs were age-residualized.

**Table 5.** Regression of DNAm at Wave 2 on Interactions between DNAm at Wave 1 and SES Covariates.

|  | PhenoAgeAccel<br>(Wave 2) |  | GrimAgeAccel (Wave 2) |  | Dunedin PACE (Wave 2) |  | Epigenetic-g (Wave 2) |  |
| --- | --- | --- | --- | --- | --- | --- | --- | --- |
| *Model 1 (n=428)* | *std. Beta* | *95% CI* | *std. Beta* | *95% CI* | *std. Beta* | *95% CI* | *std. Beta* | *95% CI* |
| MPS Wave 1 | **0.43\*** | 0.29, 0.58 | **0.52\*** | 0.39, 0.65 | **0.31\*** | 0.17, 0.45 | **0.45\*** | 0.31, 0.59 |
| Family SES | 0.06 | -0.10, 0.23 | 0.01 | -0.13, 0.15 | -0.17 | -0.33, 0.00 | 0.06 | -0.09, 0.22 |
| COVID SES Stress | -0.04 | -0.20, 0.13 | 0.04 | -0.09, 0.18 | 0.03 | -0.14, 0.20 | 0.11 | -0.05, 0.26 |
| MPS Wave 1*Family SES | 0.09 | -0.08, 0.26 | 0.12 | -0.04, 0.28 | 0.03 | -0.15, 0.21 | 0.02 | -0.13, 0.17 |
| MPS Wave 1*COVID SES Stress | 0.08 | -0.08, 0.25 | 0.08 | -0.06, 0.23 | -0.07 | -0.21, 0.06 | 0.02 | -0.13, 0.17 |
| *Model 2 (n=428)* | *std. Beta* | *95% CI* | *std. Beta* | *95% CI* | *std. Beta* | *95% CI* | *std. Beta* | *95% CI* |
| MPS Wave 1 | **0.43\*** | 0.28, 0.58 | **0.43\*** | 0.27, 0.59 | **0.29\*** | 0.15, 0.44 | **0.41\*** | 0.26, 0.56 |
| Family SES | 0.06 | -0.11, 0.22 | 0.00 | -0.15, 0.14 | -0.13 | -0.29, 0.03 | 0.04 | -0.12, 0.20 |
| COVID SES Stress | -0.04 | -0.23, 0.14 | -0.01 | -0.16, 0.14 | 0.02 | -0.15, 0.20 | 0.13 | -0.04, 0.29 |
| MPS Wave 1*Family SES | 0.07 | -0.10, 0.25 | 0.09 | -0.08, 0.25 | 0.07 | -0.11, 0.26 | 0.04 | -0.11, 0.20 |
| MPS Wave 1*COVID SES Stress | 0.09 | -0.08, 0.26 | 0.08 | -0.06, 0.23 | -0.04 | -0.17, 0.10 | 0.01 | -0.14, 0.17 |
| Assessment Lag | 0.14 | -0.04, 0.32 | 0.15 | -0.00, 0.30 | -0.03 | -0.21, 0.14 | 0.13 | -0.04, 0.29 |
| Sex (Male) | -0.27 | -0.57, 0.03 | **0.34** | 0.03, 0.64 | **-0.36** | -0.66, -0.07 | -0.13 | -0.42, 0.16 |
| Latinx | 0.00 | -0.17, 0.17 | -0.00 | -0.14, 0.14 | **0.20** | 0.03, 0.38 | -0.06 | -0.22, 0.09 |
| BlackPlus | 0.15 | -0.03, 0.32 | 0.11 | -0.04, 0.26 | 0.09 | -0.08, 0.27 | -0.09 | -0.26, 0.07 |
| AsianPlusOther | 0.06 | -0.11, 0.23 | 0.05 | -0.09, 0.20 | -0.06 | -0.23, 0.11 | -0.08 | -0.24, 0.08 |
| LatinxWhite | 0.02 | -0.16, 0.19 | 0.01 | -0.13, 0.16 | 0.14 | -0.03, 0.31 | -0.09 | -0.26, 0.07 |
| DNAm-Smoke | 0.01 | -0.15, 0.17 | 0.07 | -0.09, 0.22 | 0.02 | -0.13, 0.18 | -0.00 | -0.15, 0.15 |
| *Model 3 (N=146)* | *std. Beta* | *95% CI* | *std. Beta* | *95% CI* | *std. Beta* | *95% CI* | *std. Beta* | *95% CI* |
| MPS Wave 1 | **0.40\*** | 0.23, 0.56 | **0.43\*** | 0.26, 0.61 | **0.32\*** | 0.16, 0.47 | **0.37\*** | 0.21, 0.54 |
| Family SES | 0.07 | -0.11, 0.25 | -0.03 | -0.18, 0.13 | -0.08 | -0.26, 0.10 | 0.05 | -0.12, 0.21 |
| COVID SES Stress | -0.03 | -0.23, 0.17 | 0.01 | -0.16, 0.18 | 0.00 | -0.19, 0.20 | 0.16 | -0.02, 0.33 |
| MPS Wave 1*Family SES | 0.10 | -0.10, 0.30 | 0.09 | -0.08, 0.27 | 0.10 | -0.10, 0.30 | 0.10 | -0.07, 0.27 |
| MPS Wave 1* COVID SES Stress | 0.12 | -0.06, 0.30 | 0.08 | -0.08, 0.23 | -0.07 | -0.23, 0.08 | 0.03 | -0.14, 0.19 |
| Assessment Lag | 0.13 | -0.08, 0.34 | 0.14 | -0.04, 0.31 | 0.05 | -0.15, 0.26 | 0.08 | -0.10, 0.27 |
| Sex (Male) | -0.27 | -0.60, 0.06 | **0.35** | 0.01, 0.69 | **-0.40** | -0.73, -0.07 | -0.08 | -0.39, 0.24 |
| Latinx | -0.02 | -0.21, 0.16 | -0.03 | -0.19, 0.12 | **0.23** | 0.04, 0.42 | -0.04 | -0.21, 0.12 |
| BlackPlus | 0.11 | -0.09, 0.30 | 0.11 | -0.06, 0.28 | 0.15 | -0.05, 0.34 | -0.17 | -0.35, 0.01 |
| AsianPlusOther | 0.07 | -0.12, 0.26 | 0.07 | -0.09, 0.23 | -0.01 | -0.20, 0.18 | -0.13 | -0.30, 0.04 |
| LatinxWhite | 0.03 | -0.16, 0.21 | 0.01 | -0.15, 0.17 | 0.12 | -0.07, 0.31 | -0.05 | -0.22, 0.12 |
| DNAm-Smoke | 0.10 | -0.08, 0.29 | 0.13 | -0.05, 0.31 | 0.09 | -0.09, 0.27 | 0.02 | -0.15, 0.19 |
| BMI (Wave 1) | 0.08 | -0.08, 0.24 | 0.03 | -0.11, 0.18 | 0.02 | -0.14, 0.17 | 0.00 | -0.15, 0.15 |
| PDS (Wave 1) | 0.21 | -0.01, 0.44 | 0.18 | -0.03, 0.38 | -0.01 | -0.24, 0.21 | **0.23** | 0.02, 0.45 |
| ΔPDS | 0.11 | -0.10, 0.32 | 0.06 | -0.13, 0.26 | 0.14 | -0.07, 0.35 | 0.08 | -0.12, 0.29 |

Note. MPS = DNA-methylation profile score. CI = Confidence Interval. Nominally significant parameters at $p < 0.05$ are in bold; parameters significant after FDR correction are denoted with \*. All MPSs were age-residualized.

## Discussion

Studies in adults have identified patterns of DNAm that reflect biological aging and that are highly stable and robust predictors of future morbidity and mortality. Children are often hypothesized to be more sensitive to environmental perturbations than are adults, but this is not always the case, and basic data regarding the longitudinal stability of MPSs is lacking. In this study, we provided the first test of the longitudinal stability of aging-related MPSs in childhood and examined whether these MPSs were affected by the environmental disruptions associated with the onset of the COVID-19 pandemic.

We had two major findings. First, in contrast to adults, who show highly stable trajectories of aging-related biology (16-year, whole blood MPS test-retest correlations ranging from 0.73 to 0.93^19^), we found moderate stability of child and adolescent salivary MPSs over about four years (with test-retest correlations comparable to child personality^46^). If lower levels of stability of MPSs during development is driven by sensitivity to environmental variation over time, our findings suggest that early life may be a more sensitive period for interventions that impact DNAm. Alternatively, greater instability in MPSs in this age range might be due to variation in pubertal development^48,49^; however, aging-related MPSs were largely unrelated to either pubertal status or change in pubertal status in our sample. Which environmental changes are capable of perturbing these trajectories, however, remain unknown. Environmental and behavioral interventions including calorie restriction and smoking cessation can affect MPSs and individual DNAm sites even in adulthood, when MPSs are highly stable.^50–52^

Second, children who were exposed to the environmental disruptions and exposures of the COVID-19 pandemic did not show significant differences in their developmental trajectories of MPSs. Children who lived through the pandemic also did not show differences in mean-level change of MPSs. The COVID-19 pandemic parallels the 1918 influenza pandemic, which had long-term negative consequences for education, disability, and socioeconomic outcomes decades after the pandemic,^53^ The shock of the COVID-19 pandemic may similarly impact developmental trajectories in ways that that do not operate via the MPSs examined here or are not apparent within our study time frame.^26^

It is of note that SES did not exhibit main effects on MPSs at follow-up incremental of baseline MPS levels, nor did SES moderate the stability of MPSs, since previous work has documented associations between socioeconomic disadvantage and accelerated pace of biological aging.^15,54,55^ That SES is not associated with later MPSs above and beyond their baseline levels suggest that socioeconomic disparities in MPSs in later adolescence and adulthood may be canalized via effects operating at earlier points in development. This is supported by recent research finding that SES at birth is more strongly associated with a MPS of BMI in childhood and adolescence than concurrent measures of SES.^24^ It is possible, however, that over longer time lags, MPSs may more substantially reorder such that (main and moderating) effects of SES on stability and change become apparent.

## Limitations

There are several limitations to note. First, we use data from only two waves spanning late childhood through adolescence, and results cannot be generalized to different time lags or different developmental stages, notably infancy and early childhood.^56^ Second, the reliability of DNAm measurement varies across probes^57^, and measurement unreliability downwardly biases estimates of longitudinal stability. However, analysis of 20 technical replicates in our sample suggests that measurement reliability of the MPSs analyzed here is good. Third, our study sample differed from the discovery samples in which MPS algorithms were derived, including differences in participant demographics and sampling strategies.^58,59^

Most notably, our study was conducted with salivary DNAm samples (consisting of a mixture of ∼65% leukocytes and ∼ 35% buccal cells), whereas the DNAm samples in adult discovery and longitudinal samples were blood-based (100% leukocytes), raising the concern that the lower longitudinal stability observed here might be driven by tissue differences rather than developmental differences. While cross-tissue DNAm comparisons remain limited, particularly in pediatric samples, results from recent investigations alleviate this concern. One study of 11-year-old children found that only 3.5% of CpG sites were differently methylated between saliva and blood;^60^ another study with participants spanning ages 13 to 73 found that blood and saliva samples showed a very high genome-wide DNAm correlation (*r* = .97).^61^ Furthermore, other salivary MPSs (for BMI and epigenetic age) have demonstrated strong longitudinal stability in adolescence.^24,25^ Together these results suggest that the relative instability of aging-related MPSs observed in the current study is not an artifact of salivary DNAm measurement. Nevertheless, as saliva is more feasible to collect in large pediatric cohorts than blood, future research on how aging-related biology changes in early life will benefit from the development of MPSs specifically trained on salivary data.

## Conclusions

The current study provides evidence for moderate rank-order stability during childhood and adolescence in MPSs that have implications for long-term health and disease. A more comprehensive understanding of canalization of the methylome is needed, given that early life environments may lead to persisting molecular changes that influence morbidity and mortality much later in life. Further investigation of MPSs during early developmental years is needed to understand what is necessary to elicit meaningful change in these indices of biological aging.

## Abbreviations

MPS: Methylation profile score
DNAm: DNA-methylation
CpG: sites in which cytosine nucleotides precede a guanine nucleotide
TTP: Texas Twin Project
EWAS: epigenome-wide association studies
BMI: body mass index
FDR: false discovery rate
SES: socioeconomic status

## Supplement

**Table S1.**
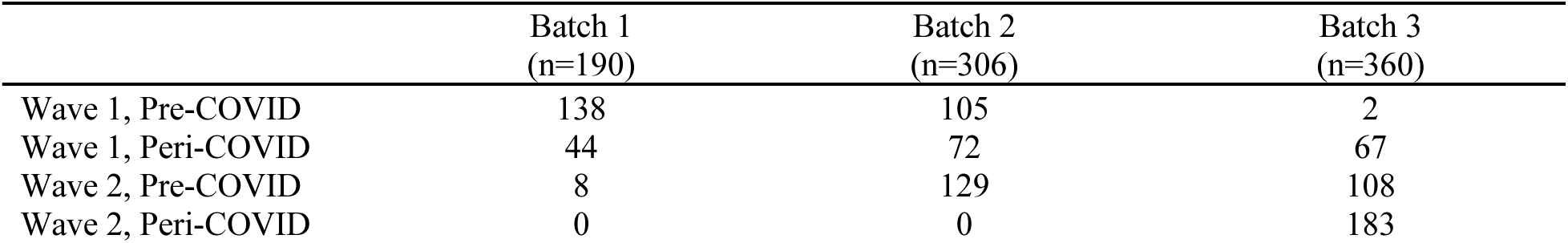
Batch, COVID-19 group, Wave Overlap of Samples.

|  | Batch 1<br>(n=190) | Batch 2<br>(n=306) | Batch 3<br>(n=360) |
| --- | --- | --- | --- |
| Wave 1, Pre-COVID | 138 | 105 | 2 |
| Wave 1, Peri-COVID | 44 | 72 | 67 |
| Wave 2, Pre-COVID | 8 | 129 | 108 |
| Wave 2, Peri-COVID | 0 | 0 | 183 |

**Table S2.**
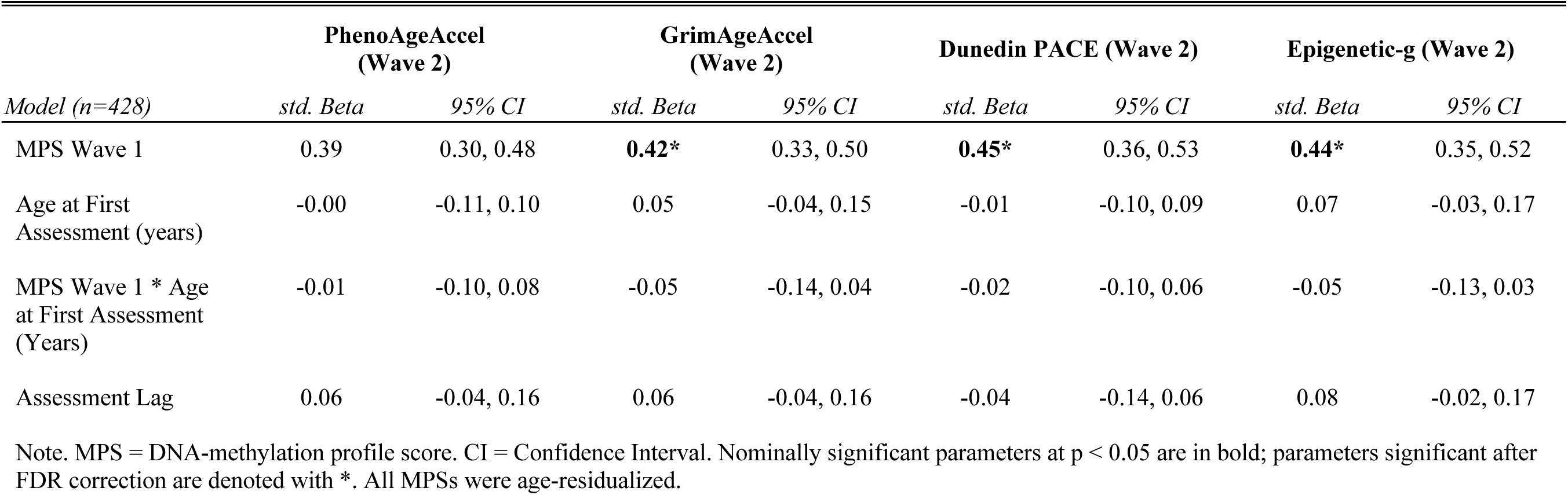
Regression of DNAm at Wave 2 on Age at First Assessment, Assessment Lag, and Interaction between DNAm at Wave 1 and Age at First Assessment (Longitudinal Stability of MPSs by Age at First Assessment)

## Notes

**Conflict of Interest Disclosures (includes financial disclosures):** The authors have indicated they have no potential conflicts of interest to disclose.

**Funding/Support:** Supported by National Institutes of Health grants R01HD083613 and R01HD092548 and the Jacobs Foundation. Drs Tucker-Drob and Harden are faculty research associates of the Population Research Center at The University of Texas at Austin, which is supported by a grant (5-R24-HD042849) from the Eunice Kennedy Shriver National Institute of Child Health and Human Development. Drs Tucker-Drob and Harden are fellows of the Jacobs Foundation.

### Competing Interest Statement

The authors have declared no competing interest.

